# Understory low compositional stability enhances functional stability in response to bull kelp removal in southern Chile

**DOI:** 10.1101/2025.01.13.632527

**Authors:** Eliseo Fica-Rojas, Daniela N. López, Alejandro Pérez-Matus, Nelson Valdivia

## Abstract

**Background and Aims.:** Ecological stability is central to understand the ability of natural systems to respond to disturbances, especially when these occur directly over important species within communities, such as foundational species. The bull kelp *Durvillaea incurvata* is common in intertidal and shallow subtidal rocky shores, supporting a diverse understory. The direct harvesting of *Durvillaea* constitute a common practice along the Chilean coast able to disrupts the abundances of the understory community and as well aggregated properties such as overall community biomass. Thus, we investigated the resistance, resilience, and recovery of the understory community of *Durvillaea* subjected to an experimental kelp removal in two sites from the temperate Chilean coast.

**Methods.:** We simulated a pulse removal of *Durvillaea* cause by direct harvesting, by removing once all the individuals within experimental plots and then we monitored the composition and biomass of the sessile and mobile understory community for 25 months.

**Key Results.:** After the disturbance we observed a rapid recovery of *Durvillaea* cover and varying stability responses of the understory community. Community composition showed a low resistance to the disturbance but returned rapidly within 5 – 7 months to pre disturbance levels (high resilience), however showed an incomplete recovery compared to controls. While community biomass displayed high resistance to the disturbance and then remained constant across time (resilience = 0), showing a complete recovery compared to control communities.

**Conclusions.:** Differing from *Durvillaea* dominated communities from other regions, our analyzed communities showed rapid recovery dynamics in compositional and functional aspects, resulting highly resilient to the type of disturbance applied. In addition, since varying stability responses reported, we encourage the use of multiple dimensions of stability to exhaustively characterize the responses of *Durvillaea* dominated communities and other kelps, when exposed to disturbances.

## Introduction

Ecological stability is central to understand how natural systems respond to disturbances (Ives & Carpenter, 2007; Donohue et al., 2013; 2016; Kéfi et al., 2019). This property corresponds to the ability of ecological systems to resist and recover from disturbances, remaining relatively unchanged over time (Harrison, 1979; Pimm, 1984; Zelnik et al., 2018). When disturbances operate directly over species with strong contribution to the community, such as foundations species, the study of ecological stability become relevant for both, fundamental and applied ecology (Cardinale et al., 2012; Hooper et al., 2012; Loreau & de Mazancourt, 2013; Battisti et al., 2016). Due its large sizes and high densities, foundation species create complex biogenic habitats and stabilize environmental variability, influencing over community structure, ecosystem functioning, and several ecosystem services provided to people (Dayton 1972; Ellison et al., 2005; Cardinale et al., 2012; Ellison 2019). In the ecological literature usually long-lived foundation species such as corals or trees, have received significant attention (Ellison 2019), while comparatively less research has been conducted to understand the stability of communities structured by short – lived and highly dynamic foundation species such as mussels or kelps (Bulleri et al. 2012, Lamy et al. 2020, Valdivia et al. 2021). Therefore, assess the impacts of disturbances over short – lived foundations species and its understory community is of widespread relevance in a context of global change.

Stability encompasses multiple dimensions (or metrics) that allow us to describe different aspect of a response to a disturbance, such as resistance, resilience, and recovery, among others (Donohue et al., 2016; Hillebrand et al., 2018; Kéfi et al., 2019). Upon a disturbance takes place, resistance involve the immediate response of the community and informs the degree to which the community remains unchanged after a disturbance; resilience is the speed or rate of return of a community to a dynamic reference state (Donohue et al., 2016; Hillebrand et al., 2018); recovery indicates the state reached by the community, relative to a reference dynamic state, occurring at the end of an observational time (Donohue et al. 2016, Hillebrand et al. 2018). These metrics have been used mainly to elucidate changes in ecological structure and functioning (Lake, 2012).

Stability dimensions can be estimated for aggregate properties performed by the whole community, such as community productivity, total cover, or biomass (functional stability; Hillebrand et al., 2018; Valdivia et al., 2021; Fica-Rojas et al., 2022) and for the combination of taxonomic identities and their abundances within the community (compositional stability; Micheli et al., 1999). Recent evidence suggests that measuring single dimensions of stability cannot reflect completely the overall stability of the system (Donohue et al., 2013, 2016; Hillebrand et al., 2018; Yang et al., 2019), because these dimensions are not necessarily correlated among them (Donohue et al., 2013; 2016; Hillebrand et al., 2018; Hillebrand & Kunze, 2020). In addition, the simultaneous assessment of compositional and functional responses allows us to identify compensation among species when one of them is lost from the community because of disturbances. Thus, estimating various stability responses of several aspects of the community will improve our mechanistic understanding of the dynamics driving community stability after disturbances (Donohue et al., 2013; 2016; Valdivia et al., 2021). However, most of studies analyze single or few stability dimensions (Donohue et al., 2016) or the response of a few species within the community (e.g., dominant species or within single trophic groups; Donohue et al., 2016; Kéfi et al., 2019).

Foundation species are numerically dominant species that form key biogenic habitats (Dayton et al., 1992; Ellison et al., 2005; Ellison, 2019). These species support the populations of many other species primarily by non-trophic and mutualistic relationships, modulating energy, nutrients, and the overall biomass of a system (Baiser et al., 2013; Ellison et al., 2005). In intertidal and subtidal habitats, kelps structure communities through the modification of abiotic and biotic conditions (Bertness et al., 1999a; 1999b; Santelices, 1990, Teagle et al., 2017). Due their large sizes and dense canopies, kelps modify local hydrodynamics, stabilize the abiotic environment, and regulate habitat availability and quality for a diverse understory community (Steneck et al., 2020; Teagle et al., 2017), mediating therefore community-level stability (Taylor & Schiel, 2005; Toohey, 2006; Stachowicz et al., 2007; Layton et al., 2019; Lamy et al., 2020; Steneck et al., 2020). The abiotic modifications produced by kelps can impact positively or negatively the understory community, but usually, seaweeds and kelp facilitate the presence of long-lived species over opportunistic species (Jenkins et al., 1999; Lilley & Schiel, 2006; Steneck et al., 2020; Vásquez et al., 2012). Therefore, disturbances that directly impact the abundance of kelps can trigger large compositional and functional stability responses in the understory.

The bull kelp *Durvillaea incurvata* (hereafter referred to as *Durvillaea*) is one of the most abundant kelps along the temperate coast of Chile between 39° S to 56° S (Santelices, 1980; Moreno, 2001). *Durvillaea* inhabits the wave-exposed low intertidal and shallow subtidal rocky shores, covering extensive areas and reaching high individual densities (up to 80 % of the shore and up to 30 individuals per m^2^; Santelices et al., 1980; Westermeier et al., 1994). As other kelps, dense and massive *Durvillaea* canopies enhance habitat availability and quality, supporting a diverse understory community (Santelices et al., 1980; Moreno, 2001; Cancino, 2016; Thomsen & South, 2019; Velásquez et al., 2020). Moreover, bull kelps provide habitat for species with ecological and economical importance, like keyhole limpets and Chilean abalone for instance (*Fissurella* spp. and *Concholepas concholepas,* respectively). Direct harvesting is a common anthropogenic disturbance affecting the populations of *Durvillaea* (Santelices et al., 1980; Bustamante & Castilla, 1990; Parra et al., 1992). After disturbances, the rapid generation times, high reproductive output, several massive recruitment events each year, and fast-growing rates of *Durvillaea* (new individuals grow up to 40 cm in a few months), may promote population recovery within one or few years (Santelices et al., 1980; Taylor & Schiel, 2005; Velásquez et al., 2020). If most understory species are adapted to the modified habitat produced by *Durvillaea*, such the rapid recovery of the foundation species should promote a rapid recovery of compositional and functional attributes in the understory.

In this study, we analyze the stability responses of understory communities to the experimental removal of the intertidal bull kelp *Durvillaea incurvata.* We analyzed the resistance, resilience, and recovery of aggregate (summed biomass across the community) and compositional properties of sessile and mobile macrobenthic understory species (invertebrates and macroalgae). In two sites of south - central Chile, we simulated a pulsed loss of bull kelp from harvesting, and then we measured species abundances in the understory for 25 months. We assessed the following two hypotheses:

Hypothesis 1: Because kelps support local biodiversity and structure local communities through facilitation, we predicted that bull kelp loss triggers significant changes in community composition and reductions in species richness and diversity. This, in turn, should be reflected in low compositional and functional resistance.

Hypothesis 2: Since rapid generation times and reproductive output promotes a rapid recovery of *Durvillaea* population, we predict that recovery patterns of the understory community will track those of the dominant kelp, resulting in a rapid compositional and functional recovery within a year or a few months after the experimental removal of *Durvillaea*.

## Material and methods

### Experimental design and sampling protocol

The study was carried out in two sites separated by approximately 40 km on the wave exposed rocky shore of south – central Chile (Figure 1): Loncoyén (39°49’19.2"S; 73°16’48.2"W) and Chaihuín (39°56′56″S; 73°34′25″W). In both sites, direct harvesting of *Durvillaea* takes place mainly during low tides of spring and summer months (Bustamante & Castilla, 1990; Velásquez et al., 2020).

**Figure 1.**
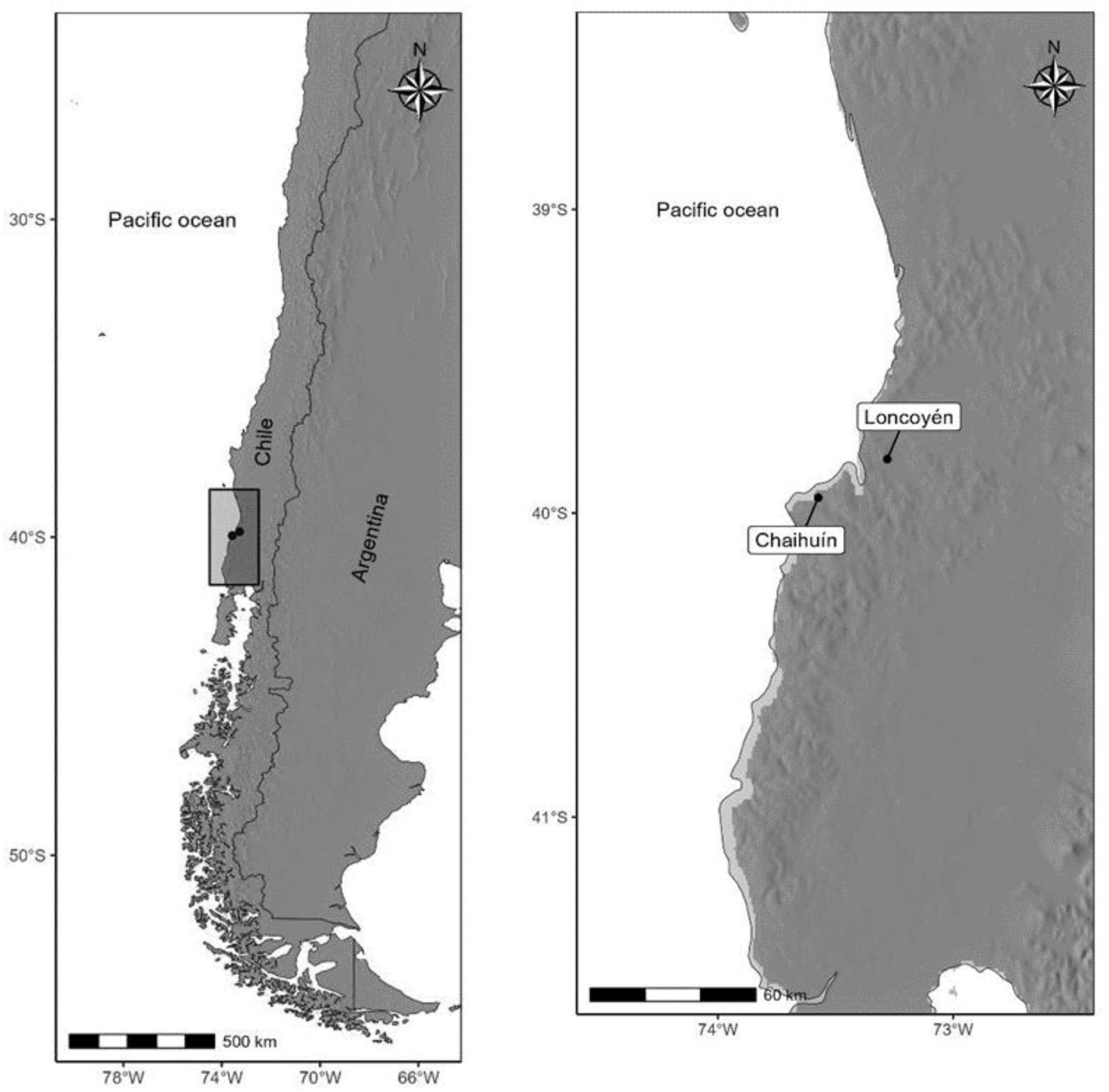
Map of the two sites, Loncoyén and Chaihuín, in south central Chile, which were used to assess the stability of *Durvillaea incurvata* dominated communities after a pulse disturbance that simulates a harvesting event.

In each site we deployed twenty 1-m^2^ plots marked with stainless-steel bolts and a numbered plastic plate. All plots were placed within areas of high cover of primary space (i.e. > 40 %) of the dominant kelp *Durvillaea*, and evenly assigned to five blocks randomly distributed along the shore (four plots per block). Each plot was haphazardly assigned to one of two treatments: either pulse disturbed or control. The disturbed treatment consisted of a single disturbance that removed once all fronds and holdfasts of *Durvillaea* from the substrate. In this way, the disturbance constituted an actual perturbation of the *Durvillaea* population (sensu Rykiel, 1985), because the disturbance generated an evident change in the abundance of this foundation species. Kelps were removed with the aid of chisels and knives. The control treatment consisted of plots in which kelp was not manipulated. Species abundances of all plots were sampled immediately before the removal of kelps, then two months after and then every three to five months until the end of the experiment. The experiment was initiated in January and October of 2020 in Chaihuín and Loncoyén respectively, and was monitored until February 2022 in Chaihuín and November 2022 in Loncoyén (25 months for each site).

At each sampling date, we used a 1-m^2^ quadrat to estimate species abundances. All seaweeds, invertebrates, and fish (c.a. > 5 mm) occurring on each experimental plot were identified in situ and the organisms were classified at the species level whenever possible. Abundance of *Durvillaea*, seaweeds and sessile invertebrates were estimated as substrate percent cover (1% resolution), while mobile invertebrates and fish were registered as density (number of individuals per m^2^). All manipulations and observations were conducted during diurnal low – tide hours.

### Estimation of understory community biomass

Percent cover and density data of the understory species were transformed to biomass (dry weight g m^2^). This was used to analyze together the aggregated biomass of mobile and sessile species, as a functional measure of the community (Fica-Rojas et al., 2022). For seaweeds and sessile invertebrates, biomass was estimated through calibration curves developed from destructive samples obtained in the field (10 to 20 replicates for the most common species identified in experimental plots). For those species whose abundances in the field were too low to generate a calibration curve, we used published linear and nonlinear models of percent cover – dry biomass for morphologically similar taxa/orders (Jung & Choi, 2022). In addition, the biomass of mobile invertebrates was estimated as the product between each species’ density (ind. m^2^) and the average individual dry weight (g) obtained from Camus et al. (2013), and from in situ samples obtained in the field (5 to 10 replicates for most common macrograzers and predators). For biomass estimation, we excluded the encrusting calcareous seaweed *Lithothamnion* sp. due to logistics constraints. Also, fish biomass was excluded from analysis due low occurrence of fish species during monitoring. Finally, community biomass was calculated for each plot as the sum of all individual’s species biomasses.

### Species richness, diversity, and structure

For each site and each sampling time, we estimated species richness (S) as the total number of taxonomic identities within each experimental plot. Then, species diversity was estimated through the Shannon diversity index (H’; Shannon 1948) following:

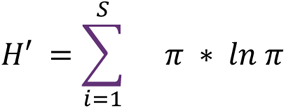

where ***π*** is the proportional abundance of species ***i.*** This metric combines both species richness and relative abundances.

Seaweeds were classified according to morphological characteristics as *Durvillaea,* kelps (others than *Durvillaea*), branched, encrusting, filamentous, foliose, leathery, and tubular seaweeds (Littler & Littler, 1984; Steneck & Dethier, 1994). The group of sessile invertebrates included species of mussels, barnacles, sponges, and ascidians; while for mobile organisms we define four groups based on relative size and trophic type, these were the mesograzers (limpets and snails, smaller than 5 cm), macrograzers (larger than 5 cm, including chitons and limpets of *Fissurella* spp.), predators (Chilean abalone, crabs, and anemones) and fish (see Supplementary table 1 for species within each group).

### Compositional and functional stability

The compositional stability of the understory community was estimated as the dissimilarity in species abundances between control and disturbed plots, calculated as Bray-Curtis distances (Bray & Curtis, 1957) in community structure between treatments. We estimated dissimilarities separately for the sessile and mobile community because these represent the resources (sessile organisms) and consumers (mobile organisms) within the community.

Functional stability was estimated from log response ratio (LRR) calculated between community biomass of control and disturbed plots as:

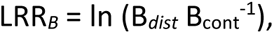

where B*_dist_* and B*_cont_* represent community biomass in the disturbed and control plots respectively. LRRs were estimated for pairs of control and disturbed plots, and for each sampling time. The pairing of plots was restricted to each block (two pairs per block).

For both compositional and functional responses, three dimensions of stability were calculated: the resistance (a), resilience (b), and recovery (c; Donohue et al., 2013; Hillebrand et al., 2018). Compositional and functional resistance were estimated as the initial dissimilarity and initial LRR, after two months since the removal of *Durvillaea*. For composition, values of *a* = 0 represent maximum resistance (no differences in composition between treatments), while values of *a* ∼ 1 present minimum resistance (maximum compositional difference between treatments). For biomass, values of ***a*** = 0 represent maximum resistance to the disturbance, while values lower or higher than 0 represent low resistance due to underperformance and overperformance relative to controls, respectively. Compositional and functional resilience (***b***) were estimated as the slopes of linear regressions of dissimilarities or LRRs over time as:

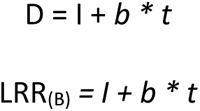

Where *I*, ***b*** and *t* represent the intercept, slope, and time respectively. Values of ***b*** = 0 represent no change over time (lack of resilience). When a < 0 (initial underperformance relative to controls), values of ***b*** < 0 implies further deviation from the control, and values of ***b*** > 0 represents more rapid recovery over time (Donohue et al., 2016; Hillebrand et al., 2018; Radchuk et al., 2019). In case that a > 0 (overperformance relative to controls), *b* < 0 represents more rapid recovery and b > 0 a further deviation of the disturbed community from the control.

Recovery (**c**) was estimated as the dissimilarity or LRR calculated at the end of the experiment (25 months after the experimental removal). For composition, *c* = 0 represents complete recovery, and *a* = 1 presents minimum recovery. For biomass, ***c*** = 0 represents complete recovery, while values lower or higher than zero represent incomplete recovery due to underperformance and overperformance relative to controls, respectively.

### Environmental monitoring

Local environmental variability during the experiment was assessed by analyzing sea surface temperature (SST) data, downloaded from the NOAA CoastWatch ERDDAP server (Simons & John 2022). Daily averaged SST data were obtained separately for both sites, from 1 x 1 km pixel obtained at 250 m from the rocky platform in which experimental plots were deployed. Seawater temperature is a useful proxy for local environmental conditions as it correlates with other environmental variables such as nutrient availability and Chlorophyll-a concentration (Nielsen & Navarrete, 2004; Witman et al., 2008) and because SST influences population growth rates of coastal marine organisms (Strathmann et al., 2002; Baldanzi et al., 2018; Suárez et al., 2020).

### Statistical analyses

We used general linear models (LMs) to assess the effect of the experimental disturbance on temporal patterns of species richness and species diversity. Also, LMs were used to estimate the effect of the experimental disturbance on the temporal patterns of percent cover of *Durvillaea* and understory seaweeds, and the density of mobile species within the understory community. Model parameters were estimated through maximum likelihood and for all the variables we used a Gaussian error distribution. Model diagnostics were checked through visual inspection of quantile-quantile plots and fitted vs. adjusted residual plots.

Non metric multidimensional scaling ordination plots (NMDS), based on Bray – Curtis dissimilarities, were used to visualize the understory sessile and mobile community structure in control and disturbed plots at each sampling time. Bray – Curtis dissimilarities were estimated separately for sessile and mobile species and the ordination was performed from several random starts until a convergent solution was reached. Permutational analysis of variance (PERMANOVA) was performed to assess the effect of the experimental removal on community structure. We used SIMPER analysis to assess species contribution to overall dissimilarities between control and disturbed plots (Clarke 1993, Clarke and Warwick 2001).

To assess if resistance, resilience, and recovery of compositional and functional responses were significantly different from reference values, we used one sample t-test. All the graphs and analyses were performed in R programming environment 4.0.2, using the “sjPlot”, “ggplot2” and “stats” packages (R Core Team, 2020). NMDS were done with the package “metaMDS” and community analyses with “vegan” (Oksanen, 2015).

## Results

### Species richness and diversity in the understory

Considering both sites, a total of 90 taxa were identified in the understory community of *Durvillaea incurvata*. These included 42 seaweeds, 31 mobile invertebrates, 15 sessile invertebrates and 2 fish species **[Supplementary table 1]**. From these species, 78 were present in Chaihuín and 53 in Loncoyén.

Prior the disturbance, species richness ranged from 12 to 15 species in both sites **[Supplementary figure 1A and 1B**]; these values were similar for controls and pulse disturbed plots (GLM – estimated [Est] between-group difference = 1.10 [S.E. = 1.72], p = 0.523, R^2^ = 0.710 and Est = -0.10 [1.29], p = 0.938, R^2^ = 0.686; for Chaihuín and Loncoyén respectively). Species richness did not differ between control and disturbed plots in Chaihuín (Est = -2.20 [2.43], p = 0.367, R^2^ = 0.710) and in Loncoyén (Est = 1.71 [1,97], p = 0.388, R^2^ = 0.686) after two months since *Durvillaea* removal, or in the following sampling times **[Supplementary figure 1A and 1B; Supplementary table 2].** Similarly, species diversity in the understory was not significantly affected by the experimental removal in both sites (**[Supplementary figure 1C and 1D]**; Est = -0.03 [0.14], p = 0.854, R^2^ = 0.130, and Est = 0.03 [0.11], p = 0.773, R^2^ = 0.149; for Chaihuín and Loncoyén respectively). For both sites and experimental treatments, species diversity varied similarly across the experiment **[Supplementary tables 2 and 3]**.

### Temporal patterns of functional groups

In Chaihuín, after two months since the removal of *Durvillaea*, the mean cover of new arrived individuals in the disturbed plots remained below the mean of controls (Figure 2A and 2B; Est = -40.20 % [9.69], p < 0.001, R^2^ = 0.328). Cover of *Durvillaea* increased steadily the following months, until reaching values similar to controls in February of 2022 (Figure 2A and 2B; Est = -0.20 % [9.69], p = 0.984, R^2^ = 0.328). Leathery seaweeds triplicated their abundances in absence of *Durvillaea* in March of 2020 (Figure 2A and 2B; Est = 10.60 % [3.94], p = 0.008, R^2^ = 0.263), but rapidly lowered when *Durvillaea* started to recover (Figure 2B). Encrusting and foliose seaweeds increased in absence of *Durvillaea* (Fig 2A and 2B), but mean levels did not differ from those of controls **[Supplementary table 4]**. Across sampling times, kelps, branched, filamentous and tubular seaweeds showed abundances below 5 % regardless of the experimental treatment (Figure 2A and 2B; **[Supplementary table 4]**). Sessile invertebrates, represented by sponges, mussels, and barnacles, ranged from 5 to 30 % across the experiment and in both treatments, reaching percent cover values greater than 25 % during August of 2020 to February of 2021 (Figure 2C and 2D). Percent cover of sessile invertebrates was not affected by the disturbance in March of 2020 (Est = 4.80% [11.38], p = 0.674, R^2^ = 0.159) or in the following sampling times **[Supplementary table 5]**. Mobile species showed similar abundance patterns for both treatments across the experiment (Figure 1E and 1F). Macrograzers and predators decreased after the removal of *Durvillaea*, while mesograzers and fish did not vary strongly across time (Figure 2E and 2F). However, for these four groups, we did not detect statistical differences between treatments after the disturbance, nor among sampling times **[Supplementary table 5]**.

**Figure 2.**
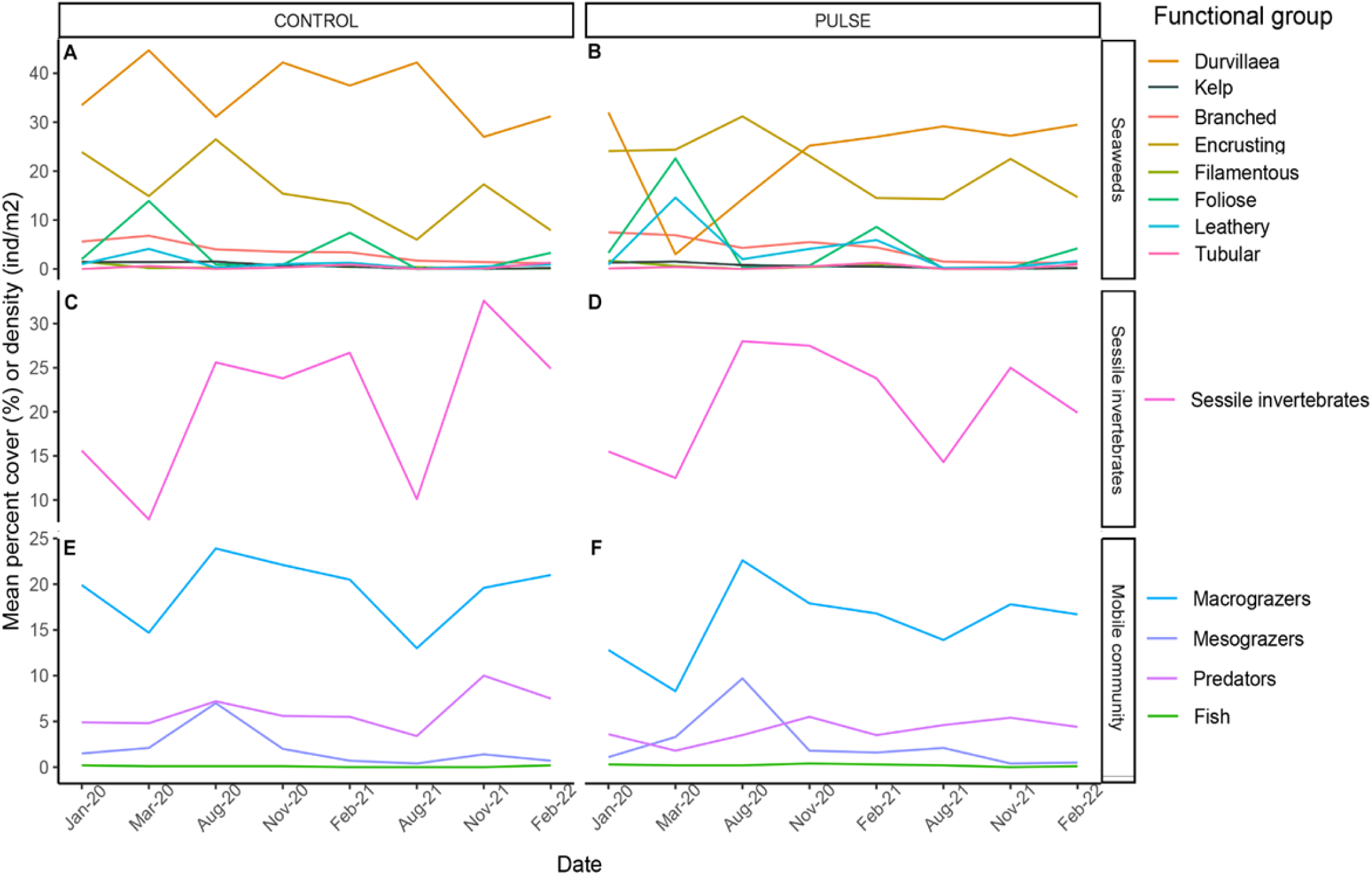
Temporal patterns of mean percent cover (seaweeds and sessile invertebrates) and density (mobile invertebrates) of the understory community of *Durvillaea* in Chaihuín, observed in control (left panels) and pulse disturbed plots (right panels, n = 10 plots per treatment). The disturbance was applied after first sampling in January of 2020. See the species assigned to each functional group in Supplementary table 1.

In Loncoyén, after two months since the removal, in December of 2020, the percent cover of *Durvillaea* differed between control and pulse disturbed plots (Figure 3A y 3B; Est = -39.46 % [10.26], p < 0.001, R^2^ = 0.492). The following sampling times the percent cover of *Durvillaea* in pulse plots remained below the mean of controls (Figure 3A and 3B; **[Supplementary table 6]**, and after 25 months, disturbed plots did not reach the same cover of controls (Est = -24.30 % [9.78], p = 0.015, R^2^ = 0.492). The cover of encrusting, branched, foliose and leathery seaweeds showed similar patterns in both treatments (Figure 3A and 3B) and were not affected by the disturbance in March of 2020 or in the following sampling times **[Supplementary table 6]**. Other kelps, filamentous and tubular seaweeds were mostly absent in both treatments across the experiment (Figure 3A and 3B; **[Supplementary table 6]**). After the disturbance and during the entire experiment, the cover of sessile species did not differ more than 1 - 2% between treatments (Figure 3C and 3D) and statistically significant between-group differences were not detected **[Supplementary table 7]**. Also, the abundances of mobile species fluctuated similarly in both treatments (Figure 3E and 3F) and no effects of the experimental removal was detected in December of 2020 or later for macrograzers, mesograzers, and predators **[Supplementary table 7]**.

**Figure 3.**
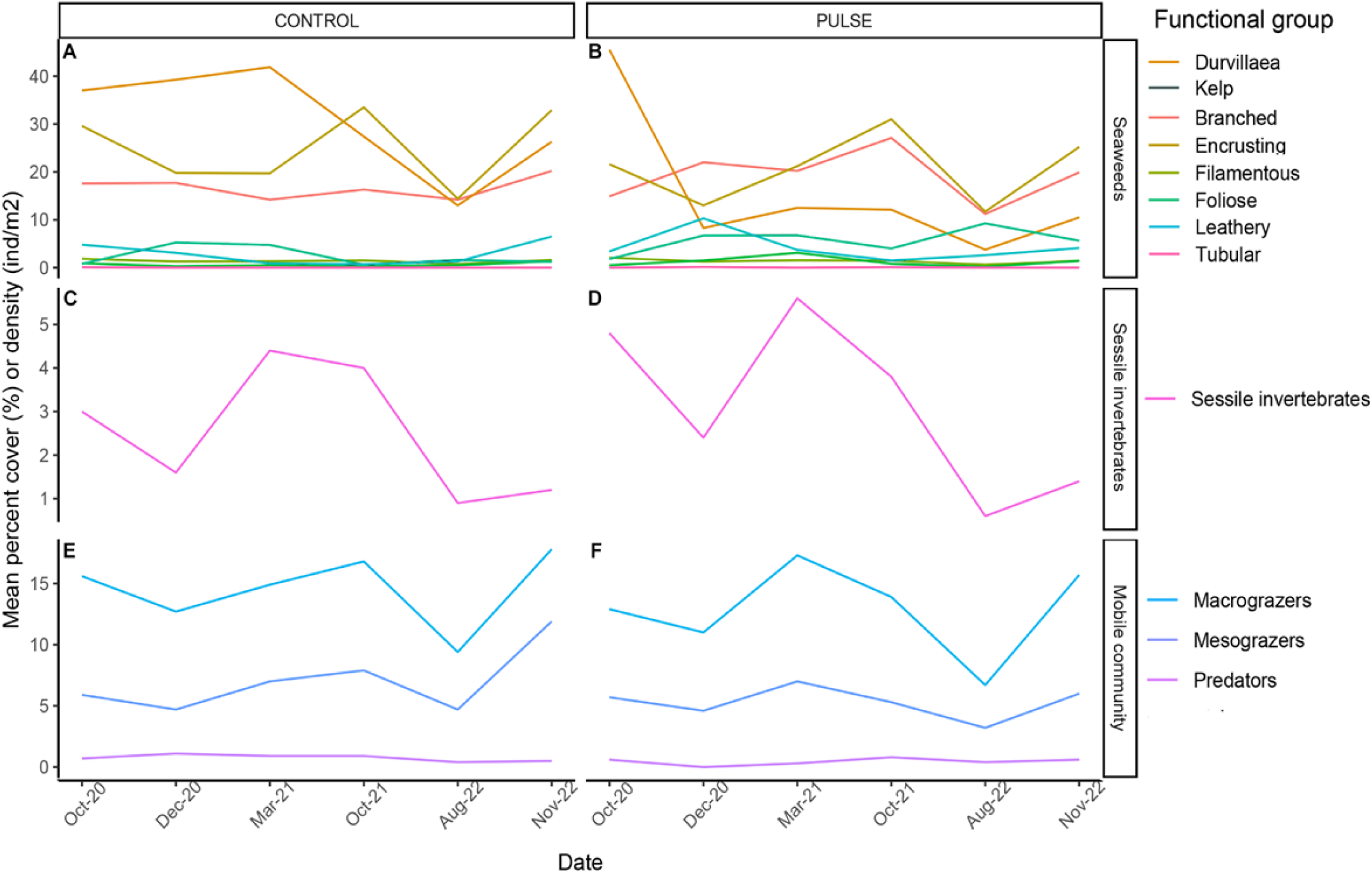
Temporal patterns of mean percent cover (seaweeds and sessile invertebrates) and density (mobile invertebrates) of the understory community of *Durvillaea* in Loncoyén, observed in control (left panels) and pulse disturbed plots (right panels; n = 10 plots per treatment). The disturbance was applied after first sampling in October of 2020. See the species assigned to each functional group in Supplementary table 1.

### Community structure and compositional stability

In Chaihuín, for the sessile community (seaweeds and invertebrates) non-metric multidimensional scaling ordination plots (NMDS) showed small temporal changes in community structure after two months since the removal of *Durvillaea* **[Supplementary figure 2A – 2H]**. Mean Bray – Curtis distances indicated an increase in between group dissimilarity after *Durvillaea* removal respect to the dissimilarity observed prior the disturbance (Figure 4A, mean between group distance [± SE] = 0.582 [0.05]) and thus, a low compositional resistance to the disturbance (Est = 0.582, t = 11.00, p < 0.001). Most of the contribution to overall between-group dissimilarities at this time were associated to the increase in disturbed plots of *Ulva* sp., *Lithothamnion* sp., Cor*allina officinallis* and *Mazzaella membranacea* **[Supplementary table 3]**. In August of 2020, 7 months after the removal, between group dissimilarity returned to pre-disturbance levels (Figure 4A, mean distance = 0.337 [0.04]). Then, dissimilarities showed small fluctuation until the end of monitoring (Figure 4A), when an incomplete compositional recovery was found (Est = 0.415, t = 9.58, p < 0.001).

**Figure 4.**
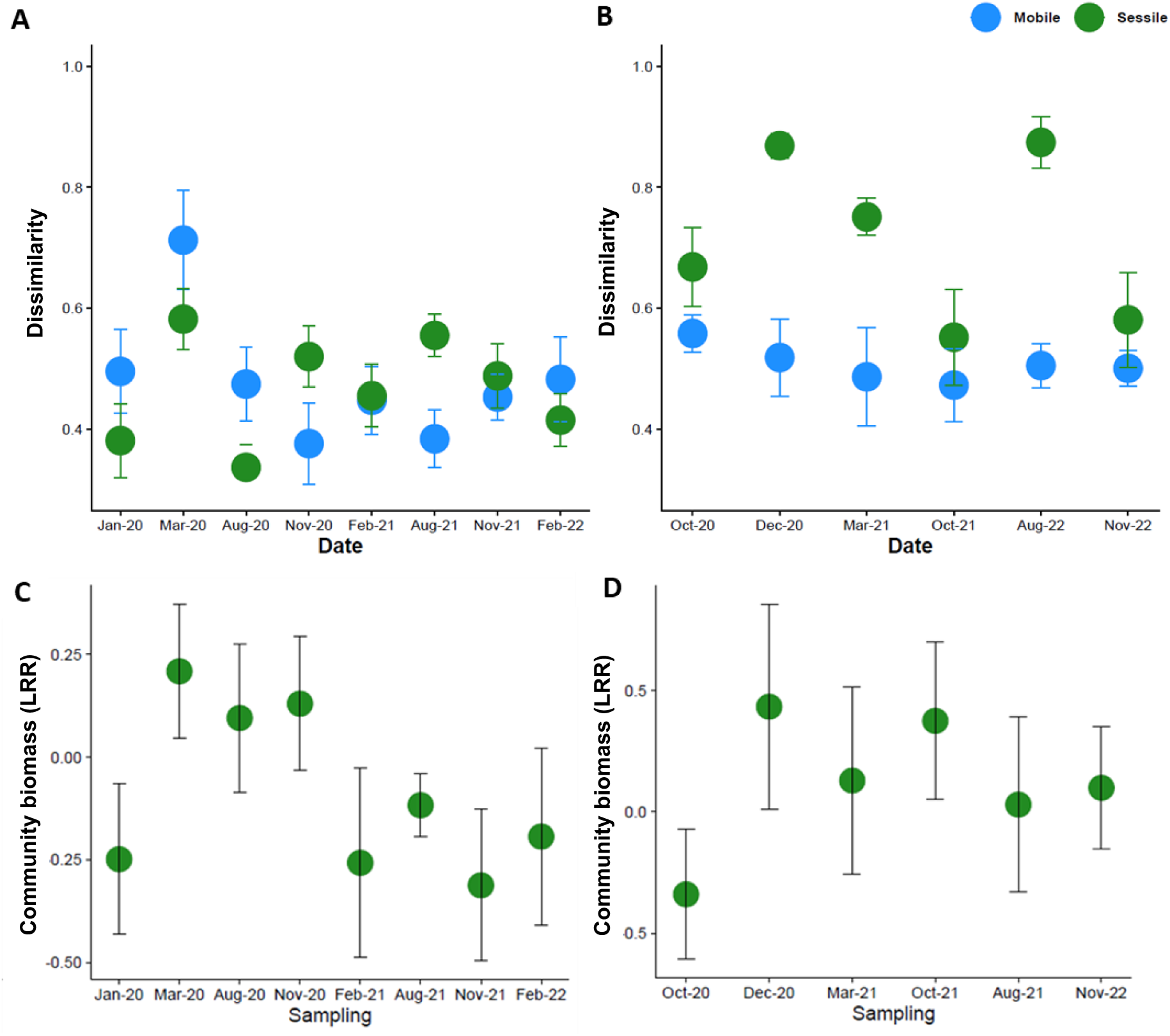
Mean between group dissimilarity of the sessile and mobile community structure, estimated as the Bray – Curtis distances between control and pulse disturbed plots, for the understory community of *Durvillaea* from Chaihuín (A) and Loncoyén (B). In bottom panels, mean log response ratio (LRR ± SE) of understory community biomass (sessile and mobile species), estimated for control and pulse disturbed plots for each sampling time for Chaihuín (C) and Loncoyén (D).

For the mobile community, NMDS evidenced that changes in community structure occurred after the removal of *Durvillaea*, but these decreased during the following sampling times **[Supplementary figure 3A – 3H]**. Bray – Curtis distances indicated an increase in between – group dissimilarity after the disturbance (Figure 4A, mean distance = 0.713 [0.08]), showing a low resistance to the disturbance (Est = 0.713, t = 8.21, p < 0.001). Dissimilarities returned to pre-disturbance levels (but not to 0) after 7 months since the disturbance, in August of 2020 (mean distance = 0.474 [0.06]) and then remained constant until the end of monitoring (Figure 3A). Accordingly, the mobile community reached an incomplete recovery at the end of the study (Est = 0.483, t = 6.91, p < 0.001). After the disturbance and at the end of monitoring, the species that contributed the most to overall dissimilarities were the macrograzers *Tonia chilensis*, *Chiton granosus*, and the predator *Concholepas concholepas* **[Supplementary tables 8 and 9]**. In addition, since the community reached pre-disturbance dissimilarities relatively fast (within less than 7 months) and then remained relatively constant, compositional resilience was not significantly different from 0 (Table 1).

**Table 1.** Results of one sample t-test for resistance, resilience, and recovery of compositional and functional responses of the understory community of *Durvillaea incurvata* in two intertidal rocky sites (Chaihuín and Loncoyén). Each dimension was compared with its reference value (µ = 0). For all measured responses, 0 represents the lack of resilience and maximum recovery. Resilience = 0, lack of. >0, high and <0, low. Recovery = 0, maximum and < 0 >, low recovery due under- or overperformance. For composition, resistance and recovery were expressed as Bray – Curtis (BC) dissimilarities between control and disturbed plots, and resilience as the temporal slope of BC. For function (community biomass), resistance and recovery were estimated as ln response ratios (LRR) between control and disturbed plots, and resilience as the slope of LRR over time.

| Site | Domain Response | Dimension | Estimate mean | t | df | p-value | CI LOW | CI HIGH |
| --- | --- | --- | --- | --- | --- | --- | --- | --- |
| Chaihuín | Compositional | Resistance sessile | 0.582 | 11.00 | 8 | **<0.001** | 0.461 | 0.704 |
|  |  | Resistance mobile | 0.713 | 8.21 | 8 | **<0.001** | 0.513 | 0.913 |
|  |  | Resilience sessile | -0.005 | -0.53 | 9 | 0.613 | -0.027 | 0.016 |
|  |  | Resilience mobile | -0.025 | -2.60 | 9 | **0.028** | -0.047 | -0.003 |
|  |  | Recovery sessile | 0.415 | 9.58 | 9 | **<0.001** | 0.317 | 0.513 |
|  |  | Recovery mobile | 0.483 | 6.91 | 9 | **<0.001** | 0.325 | 0.641 |
|  | Functional | Resistance | 0.209 | 1.27 | 9 | 0.235 | -0.163 | 0.581 |
|  |  | Resilience | -0.084 | -3.48 | 9 | **0.007** | -0.138 | -0.029 |
|  |  | Recovery | 0.194 | -0.89 | 9 | 0.393 | -0.682 | 0.294 |
| Loncoyén | Compositional | Resistance sessile | 0.869 | 32.70 | 5 | **<0.001** | 0.801 | 0.937 |
|  |  | Resistance mobile | 0.519 | 6.28 | 5 | **0.002** | 0.306 | 0.731 |
|  |  | Resilience sessile | -0.049 | -1.62 | 8 | 0.145 | -0.118 | 0.021 |
|  |  | Resilience mobile | -0.002 | -0.07 | 8 | 0.943 | -0.049 | 0.046 |
|  |  | Recovery sessile | 0.581 | 6.64 | 7 | **<0.001** | 0.374 | 0.788 |
|  |  | Recovery mobile | 0.501 | 14.90 | 7 | **<0.001** | 0.422 | 0.580 |
|  | Functional | Resistance | 0.434 | 0.85 | 6 | 0.424 | -0.805 | 1.670 |
|  |  | Resilience | -0.402 | -1.49 | 9 | 0.170 | -1.010 | 0.270 |
|  |  | Recovery | 0.099 | 0.35 | 7 | 0.736 | -0.571 | 0.770 |

In Loncoyén, NMDS showed large changes in the structure of the sessile community after the disturbance and across sampling times **[Supplementary figure 4A – 4F]**; on the contrary NMDS showed small changes in the structure of the mobile community **[Supplementary figure 5A – 5F]**. For sessile organisms, mean between – group dissimilarity increased from 0.668 [0.06] prior the disturbance, to 0.869 [0.02] after two months of *Durvillaea* removal (Figure 4B). Compositional dissimilarity decreased to pre-disturbance levels by October of 2021 (Figure 4B; mean distance = 0.552 [0.08]). Dissimilarity increased again the following sampling time (Figure 3B, mean distance = 0.874 [0.04]) and then returned to pre-disturbance level at the end of monitoring (Figure 4B, mean distance = 0.581 [0.07]). Thus, the sessile community showed a low compositional resistance to *Durvillaea* removal (Est =0.869, t = 32.70, p < 0.001), and despite pre disturbance levels were reached, an incomplete compositional recovery was found (Est = 0.581, t = 6.64, p < 0.001). The species that contributed the most to overall dissimilarities were represented by *Lithothamnion* sp., *Corallina* officinallis, *Gelidium lingulatum* and *Gelidium* sp. **[Supplementary tables 10 and 11]**.

For the mobile community, between-group dissimilarity was relatively high prior to the disturbance (Figure 4B, mean distance = 0.558 [0.03]). After the disturbance, in December of 2020, and across the experiment, between-group dissimilarity did not change from initial values observed (Figure 4B). T-test indicated that these values represented a low compositional resistance and incomplete recovery (Est = 0.519, t = 6.28, p = 0.002, and Est = 0.501, t = 14.49, p <0.001, respectively) to the disturbance; however these values were not different from initial dissimilarities. The species that most contributed to overall between-group dissimilarities were the macrograzers *Fissurella* sp. and *Chiton granosus* Supplementary tables 10 and 11). In this site, resilience values resulted equal to 0 for the sessile and the mobile community (Table 1).

### Understory community biomass and functional stability

In Chaihuín, after the experimental removal took place in March of 2020, community biomass exhibited a small and statistically non-significant overcompensation in disturbed plots with respect to controls (Figure 4C; i.e., high functional resistance, Est = 0.209, t = 1.27, p = 0.235). The LRRs of community biomass fluctuated over time around 0 until the end of the experiment (Figure 4C), showing a high functional recovery from the disturbance (Est = 0.194, t = -0.89, p = 0.393). We observed a low, but statistically significant, negative resilience since the initial small overcompensation (Est = -0.084, t = -3.48, p = 0.007). Similarly, in Loncoyén the understory community exhibited a small overcompensation of biomass after the disturbance (Figure 4D), which was statistically non-significant (strong functional resistance, Est = 0.434, t = 0.858, p = 0.424, Figure 4D). In the following sampling times LRRs fluctuated around 0 until the end of the experiment (Figure 4D), when we observed a high functional recovery (Est = 0.099, t = 0.35, p = 0.736). Because of the rapid recovery of community biomass, functional resilience resulted statistically equal to 0 in this site (Table 1).

### Environmental variability during the experiment

During the experiment, mean sea surface temperature (SST) ranged from 10 to 15° C in Chaihuín and from 12 – 18° C in Loncoyén (Supplementary figure 6). Maximum SST was ca. 16° C in Chaihuín, and ca. 20 °C in Loncoyén. Lowest temperate occurred in Chaihuín during autumn and winter of 2022 (between 10 – 11° C). During the same period, mean SST was 12.5° C in Loncoyén. Differences in temperature between both sites were larger mostly in summer months, probably due to larger inputs of cold water from the river in Chaihuín

## Discussion

In this study, the experimental removal of the bull kelp Durvillaea incurvata triggered significant and rapid responses of the understory community. After 25 months of monitoring, the understory sessile and mobile community from two sites in south-central Chile (Chaihuín and Loncoyén) showed significant compositional and functional responses to the pulse disturbance, except for compositional and functional resilience, which was equal to zero due rapid recovery dynamics of both domains. The removal initially increased compositional dissimilarity between disturbed and non-disturbed communities, and impacted the temporal patterns of branched, foliose, and encrusting seaweed abundances. Specifically, we observed a low compositional resistance of sessile and mobile communities in Chaihuín, and the sessile community in Loncoyén. A complete compositional recovery was reached within seven months since the disturbance. In contrast, community biomass at both sites showed a high functional resistance to the experimental disturbance. Then, community biomass remained relatively similar between experimental treatments across the experiment, resulting in a high functional recovery to the disturbance. Therefore, the short generation times, high reproductive output and rapid growing of bull kelps accounted for the rapid recovery of the understory community within a short period of time.

### Underlying dynamics driving community stability

Interspecific competition among understory seaweeds can regulate the recovery trajectories of disturbed kelp populations and the whole community (Wernberg et al., 2018). For example, competition for primary space (preemptive competition) mediates the recovery of intertidal communities after the experimental removal of *Ascophyllum nodosum* and *Hormosira banksii* (Lilley & Schiel, 2006; Schiel & Lilley, 2007; Phillippi et al., 2014; Schiel & Gunn, 2019). For these species, low recovery occurs because red turf-forming seaweeds inhibit the reestablishment of kelp early stages (Lilley & Schiel, 2006; Ingólfsson & Hawkins, 2008). Particularly, for *A. nodosum* the recruitment of other Fucales delays the recovery of the original dominant population and disrupts the overall recovery of the community (Ingólfsson & Hawkins, 2008). Also, turf forming seaweeds can enhance sedimentation and consequently inhibit the recruitment of kelp zygotes (Schiel & Gunn, 2019). When individuals of *Durvillaea* are removed from the intertidal, an increase in cover of *Corallina* sp. and encrusting *Lithothamnion* spp. has been shown elsewhere (Santelices, 1980; Santelices et al., 1980). In our study sites and upon canopy removal, we observed a similar positive response of seaweeds such as *Corallina officinallis, Gelidium lingulatum*, *Mazzaella membranacea* and *Ulva* sp. These seaweeds can rapidly use the released primary space and the increased light environment in the absence of *Durvillaea*, but usually are considered as ephemeral species that track environmental variation and are very sensitive to desiccation, wave forces, solar radiation and bleaching (Gómez et al., 2004; Moreno, 2001). In our disturbed plots, therefore, environmental variability might have induced mortality of understory opportunistic seaweeds, releasing available space for new recruits of *Durvillaea* and the reestablishment of the original community within a short period.

Also, competition during settlement with other kelp species such as *Lessonia* spp. or *Macrocystis pyrifera* has been suggested to delay the recovery of disturbed populations of *Durvillaea* and its associated community in central and northern Chile (Santelices et al., 1980; Westermeier et al., 1994). However, we did not find any evidence of other kelp species growing massively in the intertidal. In Loncoyén, recruits, juveniles, and adults of *Lessonia spicata* and *M. pyrifera* were observed only in low abundances in control and disturbed plots (i.e. less than 5%). In Chaihuín, we only observed recruits and juveniles of *M. pyrifera*, which were no longer than 40 cm in total length. Thus, these species may not be strong competitors of *Durvillaea* in this system, and probably their impact on community recovery dynamics was minor in our experiment.

In both sites, the removal of *Durvillaea* caused an initial colonization of opportunistic and ephemeral seaweeds that increased compositional differences between experimental treatments. However the abundance of these species returned to pre-disturbance levels when *Durvillaea* started to recover. Accordingly, the facilitation successional model may fit to these communities (Connell & Slatyer, 1977; Pulsford et al., 2014). In addition to rapid colonization occurring during succession, the increase of biomass of these species stabilized overall community biomass in the short term. This can reflect a potential compensation mechanism associated with the diversity of environmental responses of opportunistic seaweeds and increased environmental heterogeneity (Lehman & Tilman, 2000; Loreau & De Mazancourt, 2008) upon canopy removal.

Regarding mobile organisms, the presence of macro- and mesograzers such as *Chiton* spp., *Fissurella* spp., *Scurria* spp. and *Siphonaria lessonii*—whose abundances were relatively constant across the experiment in our plots—can have a significant role in the successional dynamics of intertidal communities, disrupting the colonization and establishment of seaweeds and invertebrates (Santelices, 1985; Aguilera, 2011; Aguilera & Navarrete, 2007; Camus et al., 2008, 2013). Both, macro- and mesograzers can consume microscopic and adult phases of seaweeds that were common in our disturbed plots, such as *Mazzaella laminarioides*, *Ulva* sp. and *Pyropia orbicularis* among others (Aguilera et al., 2015; Camus et al., 2008, 2013). Also, the most common chiton in our sites, *Chiton granosus*, has been described to have a positive effect in the production of bare space (Aguilera & Navarrete, 2007; Aguilera, 2011). Therefore, in the sites monitored here, both macro- and mesograzers species, might have limited a progressive increase of opportunistic and ephemeral seaweeds, indirectly enhancing the stability at the community level, by means of maintaining empty space available for *Durvillaea* recruitment and the consequent reestablishment of the original community. However, further research in this topic is necessary to better understand the role seaweeds – grazers dynamics in community stability.

### Impacts of the scale of the disturbance on community responses

In our study, compositional and functional recoveries were much faster than those reported in intertidal communities dominated by *Durvillaea* in south and central Chile (Bustamante & Castilla, 1990; Westermeier et al., 1994; Castilla et al., 2007), and in New Zealand (Taylor & Schiel, 2005; Dunmore, 2006; Schiel, 2006; Thomsen et al., 2019; Schiel et al., 2021). These differences were probably associated to the spatial extent, magnitude, duration, and frequency of the disturbance applied, which are considered as key factors influencing the dynamics of return to a reference state after a disturbance (Jones & Syms, 1998; Battisti et al., 2016; Graham et al., 2021).

The rapid responses observed in our study occurred after the application of a single removal simulating a harvesting event, which operated over 1 m^2^ replicates randomly distributed on the rocky shore. This disturbance simulates the patchy way and magnitude in which fishermen harvest the bull kelps in our study sites. In other studies, recovery of *Durvillaea* populations and the understory community has occurred after several years. For example, recovery of *Durvillaea* populations in Las Cruces was delayed due intense and repeated events of harvesting occurring at larger areas within the rocky shore (Bustamante & Castilla, 1990; Castilla et al. 2007). Other no-removal disturbances, such as frond pruning, resulted in the lack of recovery in population from southern Chile (Westermeier et al., 1994; Mansilla et al. 2007). Similarly, slow recovery trajectories of compositional and functional responses of *Durvillaea* dominated communities, occurred in New Zealand as consequence of disturbances operating at larger spatial scales, such as coastal marine heat waves, earthquakes, and coastal uplift (South et al., 2016; Thomsen et al., 2019; Schiel et al., 2021; Thomsen et al., 2021). In addition, these events were characterized by killing most of the organisms in the community, being the recovery to a reference state highly dependent on individuals that survived the disturbance or to reproductive nearby populations, as a source of kelp propagules. In our study, bull kelps were abundant in rocky platforms where we deployed experimental plots, thus we suggest that propagules coming from neighboring individuals likely recruited within a few meters and in the experimental plots, enhancing the resilience and recovery of the overall community.

## Conclusions

At the spatial scale and type of disturbance applied in our experiment, which resembled the way in which fishermen harvest bull kelps, the understory community showed a high resilience to the removal of *Durvillaea*, recovering within a period of 5 to 7 months in terms of species composition and community biomass. On the other hand, dimensions of stability such as the resistance and recovery informed different magnitudes and directions of understory community responses to the removal of *Durvillaea*. In contrast, the resilience was not a good estimator for the compositional and functional responses of these communities, due the rapid recovery observed in both sites. Therefore, incorporating multiple dimensions of stability effectively enhance our understanding of the responses of disturbed communities dominated by *Durvillaea*. In addition, we recommend that to broadly characterize the responses of *Durvillaea* dominated communities to disturbances, it is necessary to look at early and late responses, such as those informed by the resistance and recovery of the system; account with this information will properly enhance the management and conservation of these ecosystems in south – central Chile and elsewhere.

## Acknowledgements and funding

We thank all the members of the Laboratorio de Ecología Litoral (UACh) who helped during the fieldwork and laboratory activities, as well as provided constructive criticism that improved early versions of this manuscript. This study was financially supported by the FONDECYT no. 1230286 grant to N.V. and by the ANID no. 21181568 national doctoral grant to E.F.R.

## sAuthors contribution

E.F.R. and N.V. conceived the ideas for this study; E.F.R and D.L. implemented the experiment and collected the data; E.F.R, D.L. and N.V. analyzed the data; E.F.R. analyzed the data and led the writing and all authors contributed critically to the drafts and gave final approval for publication.

## Literature cited

Aguilera MA. 2011. The functional roles of herbivores in the rocky intertidal systems in Chile: A review of food preferences and consumptive effects. Revista Chilena de Historia Natural 84(2): 241–261. doi: 10.4067/S0716-078X2011000200009

Aguilera MA, Navarrete SA. 2007. Effects of *Chiton granosus* (Frembly, 1827) and other molluscan grazers on algal succession in wave exposed mid-intertidal rocky shores of central Chile. Journal of Experimental Biology and Ecology 349: 84–98. doi: 10.1016/j.jembe.2007.05.002

Aguilera MA, Valdivia N, Broitman BR. 2015. Facilitative effect of a generalist herbivore on the recovery of a perennial alga: Consequences for persistence at the edge of their geographic range. PLoS ONE, 10(12): 1–23. doi: 10.1371/journal.pone.0146069

Baiser B, Buckley HL, Gotelli NJ, Ellison AM. 2013. Predicting food-web structure with metacommunity models. Oikos 122(4): 492–506. doi: 10.1111/j.1600-0706.2012.00005.x

Baldanzi S, Storch D, Navarrete SA, Graeve M, Fernández M. 2018. Latitudinal variation in maternal investment traits of the kelp crab *Taliepus dentatus* along the coast of Chile. Marine Biology 165(2): 1–12. doi: 10.1007/s00227-018-3294-2

Battisti C, Poeta G, Fanelli G. 2016. An Introduction to Disturbance Ecology. New York City, NY: Springer International Publishing.

Bertness MD, Leonard GH, Levine JM, Bruno JF. 1999. Climate-driven interactions among rocky intertidal organisms caught between a rock and a hot place. Oecologia 120(3): 446–450. doi: 10.1007/s004420050877

Bertness MD, Leonard GH, Levine JM, Schmidt PR, Ingraham AO. 1999. Testing the Relative Contribution of Positive and Negative Interactions in Rocky Intertidal Communities. Ecology 80(8): 2711. doi: 10.2307/177252

Bray JR, Curtis JT. 1957. An Ordination of the Upland Forest Communities of Southern Wisconsin. Ecological Monographs 27(4): 325–349. doi: 10.2307/1942268

Bustamante RH, Castilla JC. 1990. Impact of human exploitation on populations of the intertidal Southern Bull-kelp *Durvillaea antarctica* (Phaeophyta, Durvilleales) in central Chile. Biological Conservation 52(3): 205–220. doi: 10.1016/0006-3207(90)90126-A

Camus PA, Daroch K, Opazo LF. 2008. Potential for omnivory and apparent intraguild predation in rocky intertidal herbivore assemblages from northern Chile. Marine Ecology Progress Series 361: 35–45. doi: 10.3354/meps07421

Camus PA, Arancibia PA, Ávila-Thieme I. 2013. A trophic characterization of intertidal consumers on Chilean rocky shores. Revista de Biología Marina y Oceanografía 48(3): 431–450. doi: 10.4067/s0718-19572013000300003

Cardinale BJ, Duffy JE, Gonzalez A, et al. 2012. Biodiversity loss and its impact on humanity. Nature 486(7401): 59–67. doi: 10.1038/nature11148

Castilla JC, Campo MA, Bustamante RH. 2007. Recovery of *Durvillaea antarctica* (Durvilleales) inside and outside Las Cruces marine reserve, Chile. Ecological Applications 17(5): 1511– 1522. doi: 10.1890/06-1285.1

Dayton PK. 1972. Toward an understanding of community resilience and the potential effects of enrichments to the benthos at McMurdo Sound, Antartica. B.C. Parker (Ed.), Proceedings of the Colloquium on Conservation Problems in Antarctica, Allen Press (1972), pp. 81–96.

Dayton PK, Tegner MJ, Parnell PE, Edwards PB. 1992. Temporal and spatial patterns of disturbance and recovery in a kelp forest community. Ecological Monographs 62(3): 421–445. doi: 10.2307/2937118

Donohue I, Petchey OL, Montoya JM, et al. 2013. On the dimensionality of ecological stability. Ecology Letters 16(4): 421–429. doi: 10.1111/ele.12086

Donohue I, Hillebrand H, Montoya JM, et al. 2016. Navigating the complexity of ecological stability. Ecology Letters 19(9): 1172–1185. doi: 10.1111/ele.12648

Dunmore R. 2006. Demography of Early Life Stages of Habitat-forming Intertidal Fucoid Algae. School of Biological Sciences, University of Canterbury, Christchurch.

Ellison AM. 2019. Foundation Species, Non-trophic Interactions, and the Value of Being Common. iScience 13: 254–268. doi: 10.1016/j.isci.2019.02.020

Ellison AM, Bank MS, Clinton BD, et al. 2005. Loss of foundation species: Consequences for the structure and dynamics of forested ecosystems. Frontiers in Ecology and the Environment 3(9): 479–486. doi: 10.1890/1540-9295(2005)003[0479:LOFSCF]2.0.CO;2

Fica-Rojas E, Catalán AM, Broitman BR, Pérez-Matus A, Valdivia N. 2022. Independent Effects of Species Removal and Asynchrony on Invariability of an Intertidal Rocky Shore Community. Frontiers in Ecology and Evolution 10: 1–14. doi: 10.3389/fevo.2022.866950

Gómez I, López-Figueroa F, Ulloa N, Morales V, Lovengreen C, Huovinen P, Hess S. 2004. Patterns of photosynthesis in 18 species of intertidal macroalgae from southern Chile. Marine Ecology Progress Series 270: 103–116. doi: 10.3354/meps270103

Graham EB, Averill C, Bond-Lamberty B, et al. 2021. Toward a Generalizable Framework of Disturbance Ecology Through Crowdsourced Science. Frontiers in Ecology and Evolution 9: 588940. doi: 10.3389/fevo.2021.588940

Harrison GW. 1979. Stability under Environmental Stress: Resistance, Resilience, Persistence, and Variability. The American Naturalist 113(5): 659–669. doi: 10.1086/283424

Hillebrand H, Kunze C. 2020. Meta-analysis on pulse disturbances reveals differences in functional and compositional recovery across ecosystems. Ecology Letters 23(3): 575–585. doi: 10.1111/ele.13457

Hillebrand H, Langenheder S, Lebret K, Lindström E, Östman Ö, Striebel M. 2018. Decomposing multiple dimensions of stability in global change experiments. Ecology Letters 21(1): 21–30. doi: 10.1111/ele.12867

Hooper DU, Adair EC, Cardinale BJ, et al. 2012. A global synthesis reveals biodiversity loss as a major driver of ecosystem change. Nature 486(7401): 105–108. doi: 10.1038/nature11118

Ingólfsson A, Hawkins SJ. 2008. Slow recovery from disturbance: a 20 year study of *Ascophyllum* canopy clearances. Journal of the Marine Biological Association of the UK 88(04): 689–691. doi: 10.1017/S0025315408001161

Ives AR, Carpenter SR. 2007. Stability and diversity of ecosystems. Science 317(5834): 58–62. doi: 10.1126/science.1133258

Jenkins SR, Hawkins SJ, Norton TA. 1999. Direct and indirect effects of a macroalgal canopy and limpet grazing in structuring a sheltered inter-tidal community. Marine Ecology Progress Series 188: 81–92. doi: 10.3354/meps188081

Jones GP, Syms C. 1998. Disturbance, habitat structure and the ecology of fishes on coral reefs. Australian Journal of Ecology 23(3): 287–297.

Jung SW, Choi CG. 2022. Estimation of Marine Macroalgal Biomass Using a Coverage Analysis. Journal of Marine Science and Engineering 10: 1676. doi: 10.3390/jmse10111676

Kéfi S, Domínguez-garcía V, Donohue I, et al. 2019. Advancing our understanding of ecological stability. Ecology Letters 22(9): 1349–1356. doi: 10.1111/ele.13340

Lamy T, Koenigs C, Holbrook SJ, Miller RJ, Stier AC, Reed DC. 2020. Foundation species promote community stability by increasing diversity in a giant kelp forest. Ecology 101(5): 1–11. doi: 10.1002/ecy.2987

Layton C, Shelamoff V, Cameron MJ, Tatsumi M, Wright JT, Johnson CR. 2019. Resilience and stability of kelp forests: The importance of patch dynamics and environment-engineer feedbacks. PLoS ONE 14(1): e0210220. doi: 10.1371/journal.pone.0210220

Lehman CL, Tilman D. 2000. Biodiversity, stability, and productivity in competitive communities. American Naturalist 156(5): 534–552. doi: 10.1086/303402

Lilley SA, Schiel DR. 2006. Community effects following the deletion of a habitat-forming alga from rocky marine shores. Oecologia 4: 672–681. doi: 10.1007/s00442-006-0411-6

Littler MM, Littler DS. 1984. Relationships between macroalgal functional form groups and substrata stability in a subtropical rocky-intertidal system. Journal of Experimental Marine Biology and Ecology 74(1): 13–34. doi: 10.1016/0022-0981(84)90035-2

Loreau M, de Mazancourt C. 2008. Species synchrony and its drivers: Neutral and nonneutral community dynamics in fluctuating environments. American Naturalist 172(2): doi: 10.1086/589746

Loreau M, de Mazancourt C. 2013. Biodiversity and ecosystem stability: A synthesis of underlying mechanisms. Ecology Letters 16: 106–115. 10.1111/ele.12073

Micheli F, Cottingham KL, Bascompte J, et al. 1999. The Dual Nature of Community Variability. Oikos 85: 161–169.

Moreno CA. 2001. Community patterns generated by human harvesting in Chilean shores: a review. Aquatic conservation: marine and freshwater ecosystems 11: 19–30.

Newman EA. 2019. Disturbance Ecology in the Anthropocene. Frontiers in Ecology and Evolution 7: 147. doi: 10.3389/fevo.2019.00147

Nielsen KJ, Navarrete SA. 2004. Mesoscale regulation comes from the bottom-up: Intertidal interactions between consumers and upwelling. Ecology Letters 7: 31–41. doi: 10.1046/j.1461-0248.2003.00542.x

Parra B, Moreno C, Westermeier R. 1992. Effect of Human Exploitation on the Intertidal Community Structure at the Valdivian Coast, Chile. In Coastal Plant Communities of Latin America. ACADEMIC PRESS, INC. 10.1016/B978-0-08-092567-7.50010-6

Phillippi A, Tran K, Perna A. 2014. Does intertidal canopy removal of *Ascophyllum nodosum* alter the community structure beneath? Journal of Experimental Marine Biology and Ecology 461: 53–60. doi: 10.1016/j.jembe.2014.07.018

Pimm SL. 1984. The complexity and stability of ecosystems. Nature 307(5949): 321–326. doi: 10.1038/307321a0

Radchuk V, Laender F, Cabral JS, et al. 2019. The dimensionality of stability depends on disturbance type. Ecology Letters 22(4): 674–684. doi: 10.1111/ele.13226

Rykiel EJ. 1985. Towards a definition of ecological disturbance. Australian Journal of Ecology 10: 361–365.

Santelices B. 1980. Phytogeographic characterization of the temperate coast of Pacific South America. Phycologia 192: 1–12. doi: 10.2216/i0031-8884-19-1-1.1

Santelices B, Castilla JC, Cancino J, Schmiede P. 1980. Comparative ecology of *Lessonia nigrescens* and *Durvillaea antarctica* (Phaeophyta) in Central Chile. Marine Biology 59: 119–132.

Santelices B. 1985. Marine herbivory studies. The South American contribution. Revista Chilena de Historia Natural 60: 153–158.

Santelices B. 1990. Patterns of organizations of intertidal and shallow subtidal vegetation in wave exposed habitats of central Chile. Hydrobiologia 92: 35–57.

Schiel DR. 2006. Rivets or bolts? When single species count in the function of temperate rocky reef communities. Journal of Experimental Marine Biology and Ecology 338(2): 233–252. doi: 10.1016/j.jembe.2006.06.023

Schiel DR, Gerrity S, Orchard S, Alestra T, Dunmore RA, Falconer T, Thomsen MS, Tait LW. 2021. Cataclysmic Disturbances to an Intertidal Ecosystem: Loss of Ecological Infrastructure Slows Recovery of Biogenic Habitats and Diversity. Frontiers in Ecology and Evolution 9: 767548. doi: 10.3389/fevo.2021.767548

Schiel DR, Gunn TD. 2019. Effects of sediment on early life history stages of habitat-dominating fucoid algae. Journal of Experimental Marine Biology and Ecology 516: 44–50. doi: 10.1016/j.jembe.2019.04.005

Simons RA, John C. 2022. ERDDAP. https://coastwatch.pfeg.noaa.gov/erddap. Monterey, CA: NOAA/NMFS/SWFSC/ERD

Sousa WP. 1984. The role of disturbance in natural communities. Annual Review of Ecology and Systematics 15: 353–391. doi: 10.1146/annurev.ecolsys.15.1.353

South PM, Lilley SA, Tait LW, Alestra T, Hickford MJH, Thomsen MS, Schiel DR. 2016. Transient effects of an invasive kelp on the community structure and primary productivity of an intertidal assemblage. Marine and Freshwater Research 67: 103–112. doi: 10.1071/MF14211

Stachowicz JJ, Bruno JF, Duffy JE. 2007. Understanding the effects of marine biodiversity on communities and ecosystems. Annual Review of Ecology, Evolution, and Systematics 38: 739– 766. doi: 10.1146/annurev.ecolsys.38.091206.095659

Steneck RS, Dethier MN. 1994. A Functional Group Approach to the Structure of Algal-Dominated Communities. Oikos 69(3): 476–498. doi: 10.2307/3545860

Steneck RS, Graham MH, Bourque BJ, Erlandson JM, Estes JA, Tegner MJ. 2002. Kelp Forest Ecosystems: Biodiversity, Stability, Resilience and Future. Environmental Conservation 29(4): 436–459. doi: 10.1017/S0376892902000322

Strathmann RR, Hughes TP, Kuris AM, Lindeman KC, Morgan SG, Pandolfi JM, Warner RR. 2002. Evolution of local recruitment and its consequences for marine populations. Bulletin of Marine Science 70: 377–396.

Suárez JL, Hansen M, Urtubia U, Lenz M, Valdivia N, Thiel M. 2020. Season-dependent effects of ocean warming on the physiological performance of a native and a non-native sea anemone. Journal of Experimental Marine Biology and Ecology 522: 151229. doi: 10.1016/j.jembe.2019.151229

Taylor DI, Schiel DR. 2005. Self-replacement and community modification by the southern bull kelp *Durvillaea antarctica*. Marine Ecology Progress Series 288: 87–102. doi: 10.3354/meps288087

Teagle H, Hawkins SJ, Moore PJ, Smale DA. 2017. The role of kelp species as biogenic habitat formers in coastal marine ecosystems. Journal of Experimental Marine Biology and Ecology, 492: 81–98. doi: 10.1016/j.jembe.2017.01.017

Thomsen MS, South PM. 2019. Communities and attachment networks associated with primary, secondary and alternative foundation species; A case study of stressed and disturbed stands of southern bull kelp. Diversity 11(4): 1–20. doi: 10.3390/d11040056

Thomsen MS, Mondardini L, Alestra T, Gerrity S, Tait L, South PM, Lilley SA, Schiel DR, Marzinelli EM. 2019. Local Extinction of Bull Kelp (*Durvillaea* spp.) Due to a Marine Heatwave. Frontiers in Marine Sciences 6: 84. doi: 10.3389/fmars.2019.00084

Thomsen MS, Mondardini L, Thoral F, Gerber D, Montie S, South PM, Tait L, Orchard S, Alestra T, Schiel DR. 2021. Cascading impacts of earthquakes and extreme heatwaves have destroyed populations of an iconic marine foundation species. Diversity and Distributions 27(12): 2369– 2383. doi: 10.1111/ddi.13407

Toohey BD. 2006. Recovery of algal assemblages from canopy disturbance: patterns and processes over a range of reef structures. Thesis

Valdivia N, Aguilera MA, Broitman BR. 2021. High Dimensionality of the Stability of a Marine Benthic Ecosystem. Frontiers in Marine Sciences 7: 569650. doi: 10.3389/fmars.2020.569650

Valdivia N, López DN, Fica-Rojas E, et al. 2021. Stability of rocky intertidal communities, in response to species removal, varies across spatial scales. Oikos 130(8): 1385–1398. doi: 10.1111/oik.08267

Vásquez JA, Piaget N, Vega JMA. 2012. The *Lessonia nigrescens* fishery in northern Chile: “how you harvest is more important than how much you harvest*.”* Journal of Applied Phycology 24: 417–426. doi: 10.1007/s10811-012-9794-4

Velásquez M, Fraser CI, Nelson WA, Tala F, Macaya EC. 2020. Concise review of the genus *Durvillaea* Bory de Saint-Vincent, 1825. Journal of Applied Phycology 32(1): 3–21. doi: 10.1007/s10811-019-01875-w

Wernberg T, Filbee-dexter K. 2019. Missing the marine forest for the trees. Marine Ecology Progress Series 612: 209–215. doi: 10.3354/meps12867

Wernberg T, Krumhansl K, Filbee-Dexter K, Pedersen MF. 2018. Status and trends for the world’s kelp forests. World Seas: An Environmental Evaluation Volume III: Ecological Issues and Environmental Impacts, January, 57–78. 10.1016/B978-0-12-805052-1.00003-6

Westermeier R, Muller DG, Gomez I, Rivera P, Wenzel H. 1994. Population biology of *Durvillaea antarctica* and *Lessonia nigrescens* (Phaeophyta) on the rocky shores of southern Chile. Marine Ecology Progress Series 110(2–3): 187–194. doi: 10.3354/meps110187

Witman JD, Cusson M, Archambault P, Pershing AJ, Mieszkowska N. 2008. The relation between productivity and species diversity in temperate-arctic marine ecosystems. Ecology 89: 66–80. doi: 10.1890/07-1201.1

Zelnik YR, Arnoldi JF, Loreau M. 2018. The Impact of Spatial and Temporal Dimensions of Disturbances on Ecosystem Stability. Frontiers in Ecology and Evolution 6: 224. doi: 10.3389/fevo.2018.00224

